# Engineering cooperative patterns in multi-species bacterial colonies

**DOI:** 10.1101/798827

**Authors:** A. I. Curatolo, N. Zhou, Y. Zhao, C. Liu, A. Daerr, J. Tailleur, J. Huang

## Abstract

Self-organization is a hallmark of all living systems [1]. In particular, coordinated cellular behavior, commonly orchestrated at the population level through reciprocal interactions among different cell species [2–4], regulates the spatial arrangement of specialized cell types to generate tissue patterning and form complex body layouts [5, 6]. The overwhelming complexity of living systems, however, makes deciphering the underlying mechanisms difficult and limits our knowledge of basic pattern-forming mechanism *in vivo* [7, 8]. A successful strategy is then to work with synthetic, engineered systems, in which cellular interactions can be more easily tailored and studied [9–13]. Here, we demonstrate a simple mechanism through which different populations of cells can self-organize in periodic patterns. Programmed population interactions are shown to lead to coordinated out-of-phase spatial oscillations of two engineered populations of *Escherichia coli*. Using a combination of experimental and theoretical approaches, we show how such patterns arise autonomously from reciprocal density-dependent activation of cellular motility between the two species, without the need of any preexisting positional or orientational cues. Moreover, by re-designing the interaction, the original out-of-phase spatial oscillation rhythm of the two populations can be accordingly turned into in-phase oscillations. The robustness and versatility of the underlying pattern-formation process suggest that it could both be generically encountered in nature, for instance in the complex bacterial ecosystems found in biofilms [14–16], and used to promote the mixing or demixing of active particles in a controlled way.

---

We engineered *Escherichia coli* AMB1655 into two genetically distinct strains so as to enhance the motility of each species when the conjugate species is present at high density. We refer to these strains as activators A and B. To do so, we control the transcription of *cheZ* to regulate the motility of cells: the deletion of *cheZ* in both strains, which is known to cause cells to tumble permanently [17], leads to non-motile strains (See supplementary Fig. 2); reintroduction of *cheZ* can then restore a motile phenotype [18–20]. To implement the motility activation, we adopted quorum sensing (QS) systems LuxI/LuxR from *Vibrio fischeri* and LasI/LasR from *Pseudomonas aeruginosa*, which are established building blocks for engineering synthetic systems with intercellular signaling and density-dependent phenotypes [21, 22]. Specifically, LasI in activator A and LuxI in activator B separately catalyze the synthesis of freely diffusing small molecules: acyl-homoserine lactone (AHL) 3oc12HSL and 3oc6HSL, respectively (Fig. 1-a). These molecules, upon binding and activating their cognate transcriptional regulator LasR (in activator B) and LuxR (in activator A) can induce gene expression [23, 24]. The resulting circuitry enables an orthogonal and reciprocal density-dependent gene activation response in both engineered strains when tested in a transwell co-culture system (Supplementary Fig.1-a, 1-b, and SI). By further placing *cheZ* under the QS-controlled promoters, each of the engineered strains transcriptionally responds to the local proliferation of its counterpart by activating motility. Consistent with the design, the impaired motility of either activator A or activator B in semi-solid agar was substantially restored by their cognate AHL signals (Supplementary Fig. 2-a and SI).

**Figure 1.**
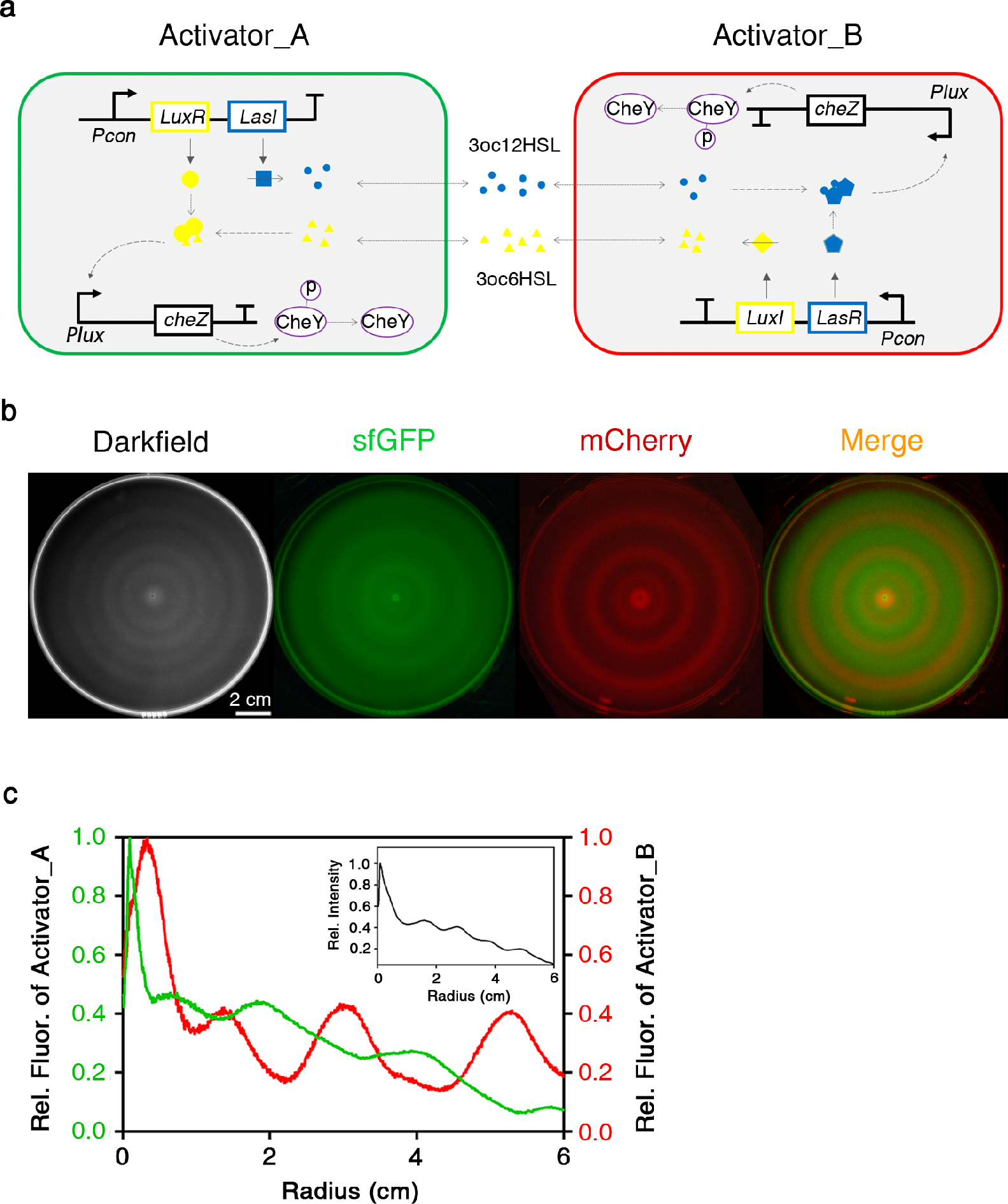
**a:** Circuit diagrams for activator strains that induce reciprocal density-dependent activation of motility. See Methods for details. **b:** Out-of-phase stripe patterns formed by the activator strains. From left to right: a darkfield snapshot of the patterns arising from the expansion of the interacting *E. coli* strains; the fluorescent signals show the presence of activators A (sfGFP) and B (mCherry). The merged fluorescent channels show the out-of-phase rythm of the density oscillations. Successive snapshots for an independent replicate are shown in Supplementary Fig. 11. **c:** Relative fluorescent intensity of activator strains as a function of the distance from the seeding center, quantified from the fluorescent images in b). Inset shows a relative density profile of the bacteria populations at varying radii for the darkfield snapshot in b).

Fig. 1-b shows a typical pattern that emerges from spatial expansion of a growing mixture of the two strains after 24 h incubation at 37°C. We mixed exponentially growing activators A and B cells with a rough 1:1 ratio and spotted the cell mixture at the center of a 14-cm Petri dish containing 27 mL 0.25% semi-solid LB1/2 agar (see SI). High density bacteria stripes separated by low density regions developed over time (Fig. 1-b, dark field, and inset of Fig. 1-c). To distinguish the two strains, each one was labeled with a spectrally distinct fluorescent protein, superfold GFP (sfGFP) and mCherry, respectively (see SI). The densities of the two activator strains (reflected by the fluorescence intensity) present out-of-phase spatial oscillations resulting in alternated fluorescent stripes, (Fig. 1-b and Fig. 1-c). Neither activator strain, when cultured alone, was capable of forming stripes (Supplementary Fig. 3-b). Similarly, noninteracting strains that express a full set of QS components but lacking density-dependent control of motility failed to form discernable patterns under the same experimental conditions, indicating that the pattern is not a result of random growth fluctuation of the engineered strains along the expansion front [25] (Supplementary Fig. 3-a and 3-b).

To further explore the self-organization capabilities offered by interspecies motility control, we engineered a second pair of strains, named inhibitors A and B, respectively, so as to negatively correlate the motility of each strain to the density of the conjugate species. To do so, we introduced a genetic inverter mediated by a repressor protein CI to reversely correlate the QS-controlled *cheZ* expression with the AHL signals (Fig. 2-a and Supplementary Fig. 1-c), hence favouring tumbling over running. Consequently, the motility of each inhibitor strain in semi-solid agar was significantly suppressed by their cognate AHL signals (Supplementary Fig. 2-b). The mixture of inhibitor A and inhibitor B, when spotted at the center of the semi-solid agar, also autonomously developed periodical stripes (Fig. 2-b, dark field and Fig. 2-c, inset). This time, however, the spatial distribution of the two strains showed in-phase oscillations resulting in colocalized fluorescent stripes, instead of the demixing observed for the activator strains (Fig.2-a and Fig.2-c). Note that, again, neither strain alone could exhibit any pattern (Supplementary Fig. 3-c).

**Figure 2.**
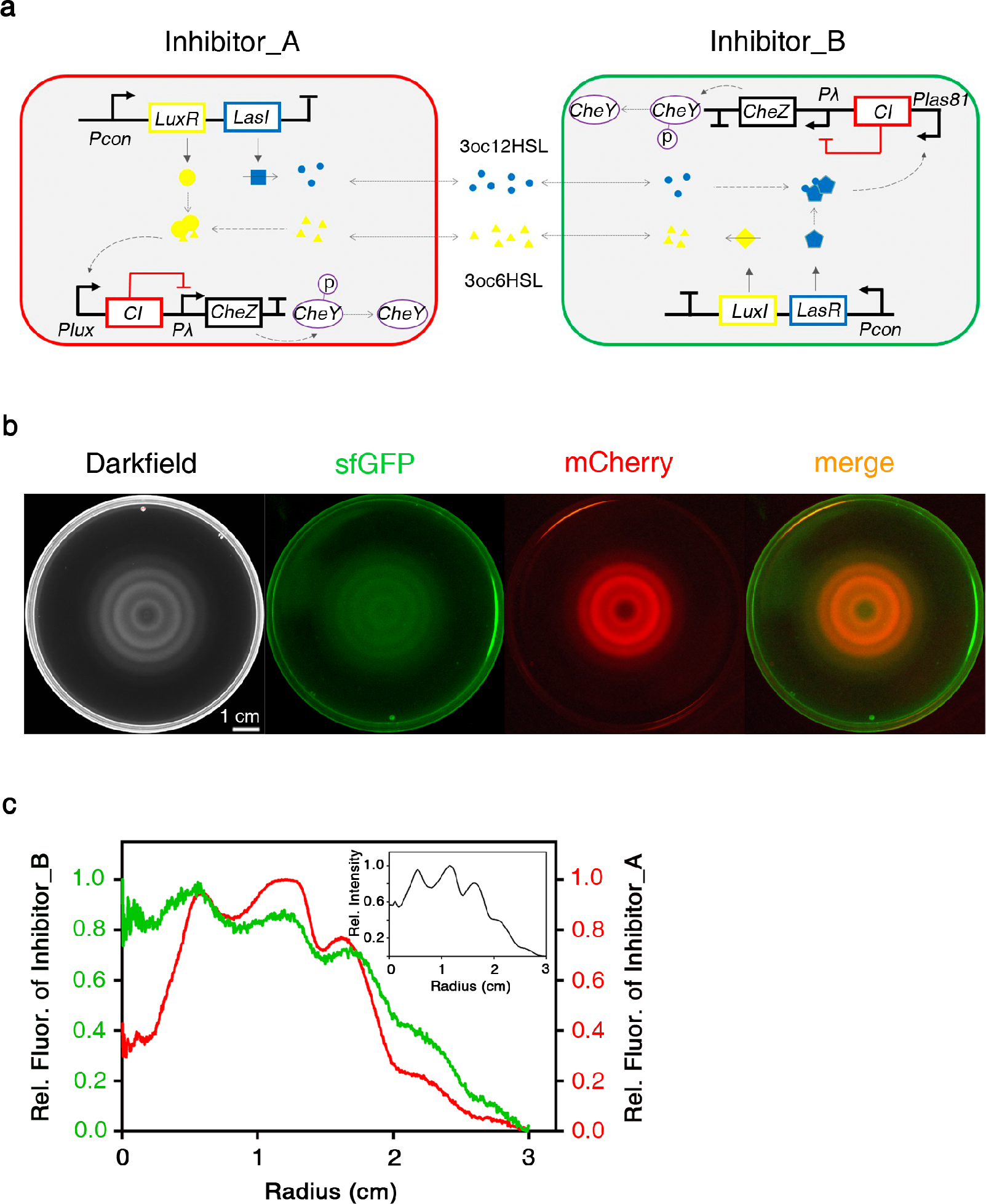
**a:** Circuit diagrams for inhibitor strains that induce reciprocal suppression of motility. See Methods for details. **b:** In-phase stripe patterns formed by the inhibitor strains. From left to right: a darkfield snapshot of the pattern resulting from the expansion of the interacting *E. coli* strains; the fluorescent signals show the presence of inhibitor A (mCherry) and B (sfGFP). The merged fluorescent channels show the in-phase rythm of the density oscillations. Successive snapshots for an independent replicate are shown in Supplementary Fig. 12. **c:** Relative fluorescent intensity of inhibitor strains as a function of the distance from the seeding center, quantified from the images in b). Inset shows a relative density profile of the bacteria populations at varying radii for the dark field snapshot in b).

To decipher the mechanisms at play in our experiments, we introduced an agent-based mathematical model comprising two types of bacteria, labeled A and B, whose dynamics stochastically alternate between runs in straight lines and tumbling phases (See Methods). Inspired by statistical physics, we follow a parsimonious approach and disregard the complexity underlying the bacterial interactions to focus on their consequences: the duration of a bacterium tumbling phase is assumed to depend on the local density of the conjugate species. The typical scales of bacterial swimming dynamics are seconds and micrometers whereas the patterns emerge on much larger scales (hours and centimeters) which are out of reach of microscopic simulations. We thus derive explicitly a coarse-grained dynamics for the density fields of each species, *ρ*_*A*_(r) and *ρ*_*B*_(r), which are shown in SI to be given by

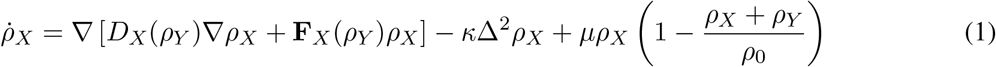

with (*X*, *Y*) describing either (*A*,*B*) or (*B*, *A*) and

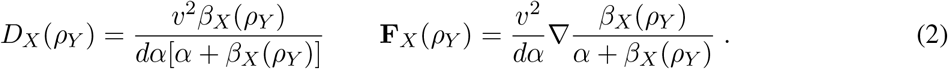

In Eq. (2), *d* is the number of spatial dimensions, *v* is the swimming speed of each species, *α* is the “tumbling rate” at which a bacterium stops swimming to start tumbling, and *β* is the “swimming rate” at which a bacterium resumes swimming in a new, random direction. For simplicity, speed and tumbling rates are assumed constant and equal for the two species, whereas the swimming rate of species *X* depends on the density *ρ*_*Y*_ of species *Y*. The last term in Eq. (1) is a logistic term that describes the population dynamics of the two species which compete for the same nutrients [7]. Finally, −*κ*Δ^2^*ρ*_*X*_ is a phenomenological regularization term whose sole role is to smooth short-range fluctuations of the density profiles. The results of simulations of Eqs. (1) with mutual activation or inhibition of the bacterial motilities, i.e. increasing and decreasing functions *β*_*X*_(*ρ*_*Y*_), are shown in Fig. 3. They perfectly correlate with our experimental results, showing that our simplified model is sufficient to account for both pattern forming processes, highlighting a single core mechanism: a density-dependent reciprocal enhancement or decrease of motility.

**Figure 3.**
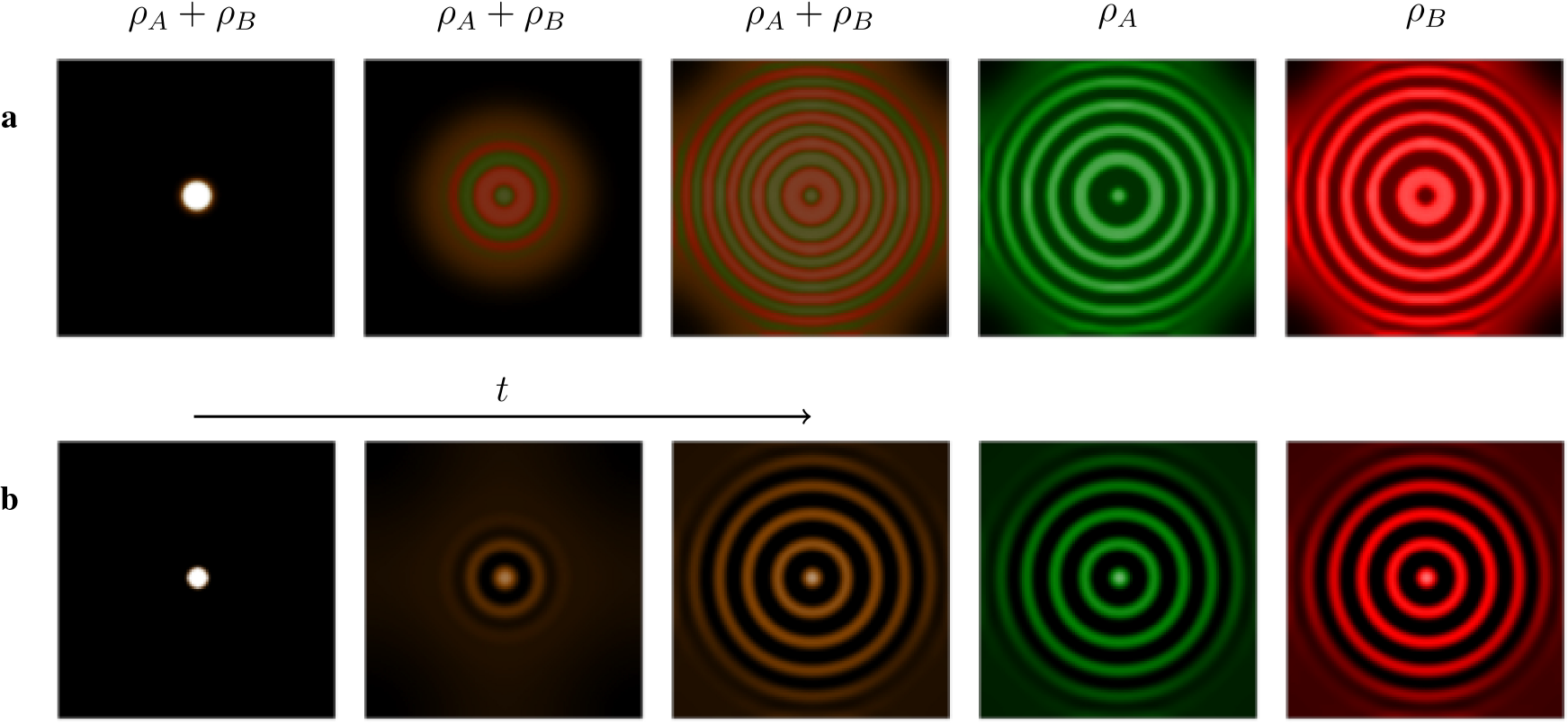
Successive snapshots of simulations of Eq. (1) starting from a central inoculum for activators (top row, left to center) and inhibitors (bottom row, left to center). The snapshots are obtained by merging a green channel representing the density of species A and a red channel representing the density of species B. At the right end of each row, the separate channels for species A and B are shown separately at a time corresponding to the central snapshot. **Row a:** Simulation parameters: 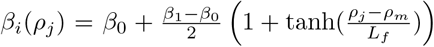, *β*_0_ = 0.5, *β*_1_ = 5, *ρ*_*m*_ = 9, *L*_*f*_ = 3, *α* = 3, *v* = 2.5, *κ* = 0:01, *μ* = 0:1, *ρ*_0_ = 16. Box size *L* = 40. Simulation times are *t* = 0, *t* = 50 and *t* = 100 from a Gaussian initial conditions with 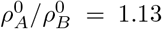. **Row b:** Simulation parameters: 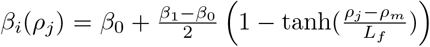, *β*_0_ = 0:5, *β*_1_ = 7, *ρ*_*m*_ = 9, *L*_*f*_ = 3, *α* = 1, *v* = 1:5, *κ* = 0:1, *μ* = 0:1, *ρ*_0_ = 24. Box size *L* = 40. Simulation times are *t* = 0, *t* = 75, *t* = 85 from a Gaussian initial conditions with 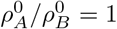.

A linear stability analysis of homogeneous density profiles 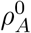 and 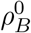, carried out in SI, sheds light on the origin of the two types of patterns observed in the experiments. The density fluxes *J*_*X*_ = −*D*_*X*_(*ρ*_*Y*_)Δ*ρ*_*X*_ − F_*X*_(*ρ*_*Y*_)*ρ*_*X*_ lead to an instability as soon as 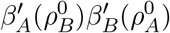 is sufficiently large. This may happen when 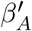 and 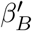 are both positive, which corresponds to a reciprocal activation of the motilities. An initial instability 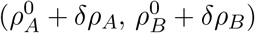 with density modulations 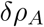 and 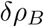 of opposite signs is then amplified exponentially, leading to the demixing of the two species. Conversely, the instability can also happen when 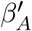 and 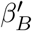 are both negative, which corresponds to mutual inhibitions of the motilities. This time, the fastest growing fluctuations correspond to 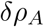 and 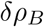 of the same sign, hence leading to the colocalization of the two species. The patterns then emerge in the wake of a travelling density wave resulting from the interplay between cell motility and population dynamics [7]. Their sizes are selected from the competition between the instability described above and the population dynamics which promote uniform densities such that *ρ*_*A*_ + *ρ*_*B*_ = *ρ*_0_ [26].

Note that while our experiments show the control of tumbling duration to be an effective lever to control cellular motility, our model predicts more general pathways: density-dependent swimming speed or tumbling rate indeed lead to similar patterns in silico (See SI). Furthermore, this mechanism appears very robust to microscopic details: comparing experimental and theoretical results shows that the experimental details ignored in the model, from the presence of the agar gel to the actual dynamics of AHL molecules or nutrient sources, indeed do not seem to strongly alter the predicted patterns.

The reciprocal control of motility is thus a versatile self-organization mechanism in bacterial colonies with two interacting species, leading to spatial structures with either aggregation or demixing between the two strains. Let us now show that this mechanism can also be used to induce the self-organization of more than two cellular species, for instance by generating pre-patterns [7]. We considered the two inhibitor strains A and B described above and designed two sensor strains, C and D, whose motility were activated or inhibited by the 3oc6HSL signal produced by inhibitor B, respectively (See SI and Supplementary Fig. 4a-c). We examined the pattern formation of the three species systems (A,B,C) or (A,B,D), containing very low fractions (1%) of the sensor strains. The cell mixture was deposited along a thin line in the middle of a semi-solid agar plate of rectangular geometry (see SI) and the patterns were recorded after 20 hours incubation at 37°C. Both experiments generated periodical patterns of alternating high- and low-density stripes, similar to the patterns arising from the interactions between inhibitors A and B (Fig.4, dark field and supplementary Fig. 4-d). We further investigated the spatial distribution of the three strains according to their distinct fluorescence labeling (inhibitor strains with sfGFP and sensor strains with mCherry fluorescence). Following the general rule that self-propelled particles accumulate where they move slower [27], sensor C, whose motility is repressed by inhibitor B, colocalized with the high-density stripes. Conversely, sensor D, whose motility is activated by inhibitor B exclusively aggregated in the low density regions (Fig.4).

**Figure 4.**
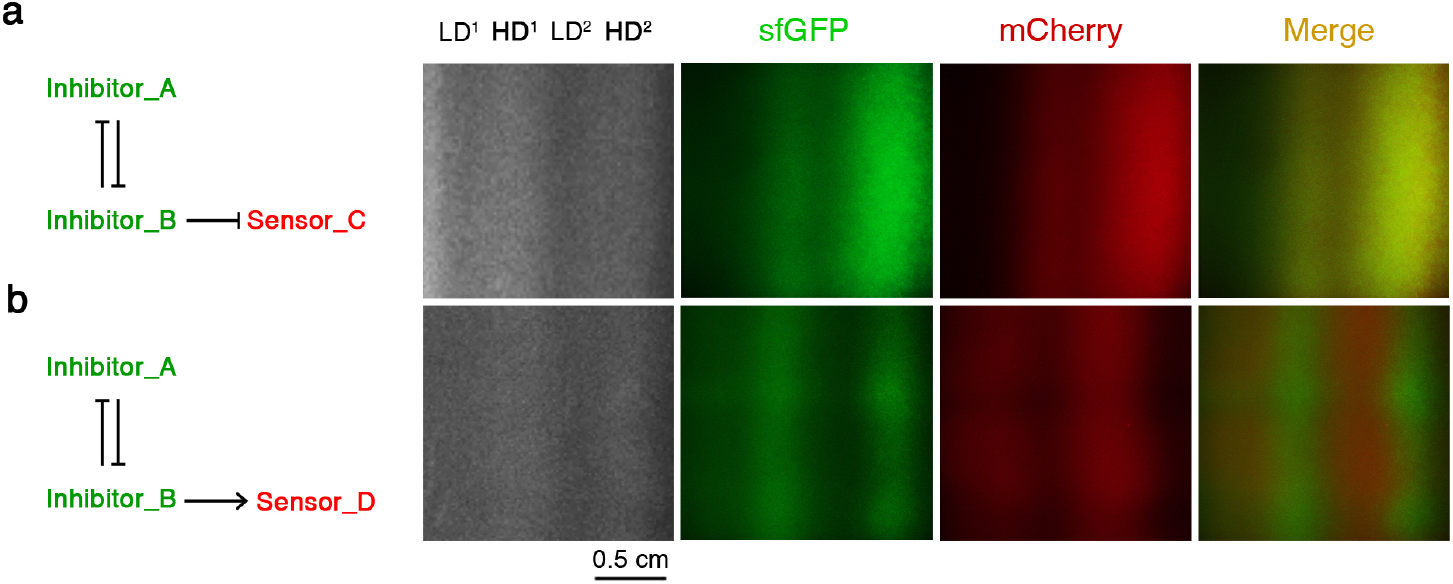
Self-organization of three-species bacterial colonies. Inhibitors A and B were mixed with a low fraction (1%) of exponentially growing sensors C or D, respectively. The three species mixture was linearly inoculated in 0.25% semi-solid LB^1/2^ agar plate and incubated at 37°C. The left panel of **a** and **b** are schematic representations of the interaction loops among the three engineered strains (see Methods and Supplementary Fig. 4-a and 4-b for details). Right panel: darkfield snapshots of stripe patterns (first column) formed by the three species systems. LD and HD indicate low- and high-density stripes, respectively. The fluorescent channels allow to distinguish the inhibitor strains A and B (sfGFP) and the sensors C and D (mCherry).

Our experimental results demonstrate that multi-species bacterial colonies are capable of self-organizing in complex macroscopic structures when the motility of each species is controlled by the density of a conjugate species. Contrary to established pattern-formation mechanisms based on chemotactic [28] or run-and-chase [3, 4] interactions, the mechanism we report does not require any directionality of the microscopic interactions: the effective attraction or repulsion between the strains only emerges at a collective level. The ensuing robustness—no microscopic chemical gradients need to be maintained for the interactions to emerge—as well as the diversity of motility control pathways should make this mechanism particularly relevant for complex bacterial ecosystems such as those found in biofilms [14–16] and porous media [29].

## Acknowledgments

We thank Mike Cates, Hugues Chaté, Wei Huang, Xiongfei Fu and Terry Hwa for discussions. AC acknowledges a doctoral fellowship from DIM ISC. AC, AD, JT, JH and YZ acknowledge support from ANR/RGC grants Bactterns & A-HKU712/14. JH acknowledges support from a National Basic Research Program of China (973 Program, 2014CB745200) from the Ministry of Science and Technology of PRC, and by Shenzhen Peacock project (KQTD2015033117210153); the Shenzhen Science and Technology Innovation Committee Basic Science Research Grant (JCYJ20150629151046896 and JCYJ20170413154523577). JCYJ20170413154523577).

## Author Contributions

AC NZ YZ AD JH JT conceived the project. NZ and JD conceived and realized the biology experiments. AC and JT conceived and realized the theoretical work. YZ and AD built the setup and realized the measurements. AC NZ YZ AD JH JT wrote the paper. CL was involved in application for grant support of the project.

## Correspondence

Correspondence and requests for materials should be addressed to Jiandong Huang and Julien Tailleur

## Methods

No statistical methods were used to predetermine sample size.

## APPENDIX A: *E. coli* strains

*E. coli* strain K12 AMB1655 Δ*lacIZYA* Δ*cheZ*, also designated as CL1, was used throughout the study to harbor genetic circuits or control plasmids unless otherwise stated. To build CL1, *Lac* operon and *cheZ* were seamlessly deleted via lambda-Red mediated homologous recombination from a parental motile *E. coli* strain K12 AMB1655 (a kind gift from Prof. Antoine Danchin) as previously described [20]. To construct noninteracting strains in Supplementary Fig. 3, *E. coli* strain K12 AMB1655 Δ*lacIZYA* with a chromosomal copy of *cheZ* was used.

## APPENDIX B: Quorum-sensing circuits

Plasmid constructions are described in SI.

## Activator strains

In both activator strains, QS receptors and biosynthetic enzymes are expressed bicistronically from a constitutive promoter (see SI). LasI in activator A catalyzes the synthesis of a membrane-diffusible small molecule 3oc12HSL (blue filled circles in Fig. 1-a). At high local 3oc12HSL concentration (reflecting high density of activator A), intracellular 3oc12HSL binds with LasR in activator B to activate the expression of CheZ. Reintroducing CheZ facilitates the dephosphorylation of CheY-phosphate (CheY-P). Similarly, LuxI in activator B catalyzes the production of 3oc6HSL (yellow filled triangles). High local 3oc6HSL concentration (reflecting high density of activator B) drives the binding of intracellular 3oc6HSL with LuxR and activates the expression of CheZ in activator A. The two activators constitutively express fluorescent proteins sfGFP and mCherry, respectively.

## Inhibitor strains

The circuitries of inhibitor A and B adopts similar QS components depicted in Fig. 2-a. To fulfill the design of mutual suppression of motility, CheZ is constitutively expressed from λ promoter (P_λ_) that can be repressed by CI repressor. A NOT gate is implemented by placing CI under the control of P_*lux*_ and P_*las81*_, a mutated Plux with stronger binding affinity with LasR [30], such that high intracellular AHLs (reflecting high cell densities) activate CI expression and in turn shut off the *cheZ* transcription. The two inhibitors constitutively express fluorescent markers mCherry and sfGFP respectively.

## APPENDIX C: Medium

The liquid medium used for bulk culture of bacteria strains is half (1/2 ×) standard Luria–Bertani (LB) medium containing 2.5 g Yeast extract, 5 g Bacto Tryptone, 5 g NaCl per liter, buffered by 0.1MHEPES (pH 7.4) unless otherwise stated. 50 *μ*g/ml kanamycin and 25 *μ*g/ml chloramphenicol were added to maintain the plasmids. Semi-solid LB^½^ agar was prepared by adding Bacto agar (BD, w/v=0.25%) into the liquid medium stated above. The mixture was autoclaved, gently mixed by shaking and stored in a 55°C water bath. Before using, 50 *μ*g/ml kanamycin, 25 *μ*g/ml chloramphenicol, 3oc6HSL (Sigma, when applicable) or 3oc12HSL (Santa Cruz, when applicable) were added to the molten agar. After vigorous mixing, 10 mL of above medium was poured into an 8.5 cm-diameter Petri dish (BD Falcon) for pattern formation of inhibitor strains; 27 mL for 14 cm-diameter Petri dish for pattern formation of activator strains. The agar was left at room temperature for at least one hour for full solidification.

## APPENDIX D: General protocols for pattern formation experiments

1. Interacting strains harboring full circuits were respectively streaked on a 1.5%(w/v) LB agar plates supplemented with 50 *μ*g/ml kanamycin and 25 *μ*g/ml chloramphenicol from corresponding glycerol stocks and incubated at 37°C overnight;
2. A single colony of each strain was inoculated into 3 mL LB medium with appropriate antibiotics and grown at 37°C with constant shaking (220 rpm) overnight;
3. The overnight culture of each strain was inoculated 1:100 into 3 mL fresh LB medium until midlog phase (OD_600_~0.4-0.6, measured by Cary Bio300 UV-Vis Spectrophotometer, Varian) at 37°C with constant shaking (220 rpm). Cells were then inoculated 1:200 into 3 mL fresh LB media with appropriate antibiotics until the cell density of each strain reaches approximately the same OD_600_ of 0.1-0.2;
4. The interacting strains were mixed using a rough 1:1 (v/v) ratio and a suspension of the mixture (2 *μ*L) was spotted at the center of a semisolid agar plate prepared on the same day;
5. The semi-solid agar plate was then moved to 37°C warm room for overnight culture (typically over 20 hours).

## APPENDIX E: Pattern formation of the three species systems in a rectangular geometry

Sensor strains were mixed in a low fraction of 1% with both inhibitors A and B. The mixture was then linearly inoculated by gently touching the semi-solid agar plate with the edge of a rectangular coverslip which was previously wetted by a liquid culture of the mixture. The plates were then left steady in 37°C warm room for overnight culture.

## APPENDIX F: Fluorescence imaging

To image the fluorescence signal of the whole petri-dish, we built the setup whose design is shown in Supplementary Fig. 9. The light from a high power white LED is collimated and filtered by a band-pass excitation filter, and then goes through the diffusor to produce homogeneous illumination for imaging. A convex lens produces an image of this diffusor in the sample plane. To minimize the amount of excitation light entering the camera, we tilt the lighting system, and use a Fresnel lens close to the sample to refocus the excitation light away from the camera. The emission filter is mounted in front of the camera (Canon EOS 700D).

To measure two different fluorescences, we built two setups in parallel that are equiped with different filter sets (Edmund Optics). For the green channel, the excitation filter is a 466±20 nm bandpass, and the emission filter is of 510±10 nm. For the red channel, the bandpass excitation filter is of 586±10 nm, and the emission filter is of 615±10 nm.

The images are taken successively using the two setups at the end of the overnight culture.

## APPENDIX G: Image acquisition and analysis

Darkfield images of the semi-solid agar plates were taken by a ChemiDoc Imaging system (Bio-Rad) equipped with a cooled CCD camera. Fluorescent images of the patterns with a circular geometry were taken by a Canon EOS 700D with an ISO setting of 800. For patterns formed by activator strains in Fig. 1 of the main text, the exposure time for GFP and mCherry channels are 1.6 s and 3.2 s respectively. For patterns formed by inhibitor strains in Fig. 2 of the main text, the exposure time for GFP and mCherry channels are 6 s and 3.2 s respectively. Fluorescent images of patterns formed by the three species systems in a rectangular geometry were taken by Leica MZ10F Fluorescence Stereomicroscopy with an exposure time of 1.2 s and gain of 1.5 for both GFP and mCherry channels. The brightness and contrasts of the images were adjusted linearly for optimal display while preserving the relative signal intensity for each image. Relative intensity analysis of the images was done using ImageJ. For patterns with circular geometry, intensity profiles (both darkfield images and fluorescent images) were calculated using the plugin “Radial Profile Angle” as the average pixel intensity values over a 2*π* angle at given radius from the seeding center. Relative values are obstained by normalizing the density profiles by their maximum and minimum.

## APPENDIX H: Theory

## 1. The model

We consider an agent-based microscopic model comprising particles of species A and B moving in two spatial dimensions (we neglect the motion of the bacteria along the vertical dimension). Particles can be either in a running state or in a *tumbling* state. In the running state, particles persistently move in a fixed direction at speed *v*_*A*_ or *v*_*B*_ depending on their species. They switch to a tumbling state at tumbling rate *α*_*A*_ or *α*_*B*_. In the tumbling state, they are motionless, until a running event, occurring at swimming rate *β*_*A*_ or *β*_*B*_, makes the cell resume swimming in a new, random direction. In our experiments, the motility parameters of bacteria of species A or B have a complex dependence—mediated by the AHL molecules and hence non-local in space and time—on the density fields of the conjugate species. As a first minimal approach, we assume interactions to be instantaneous and local. The dynamics of a particle of species A at position r is then characterized by functions *v*_*A*_(*ρ*_*B*_(r)), *α*_*A*_(*ρ*_*B*_(r)) and *β*_*A*_(*ρ*_*B*_(r)), and conversely for species B, where we have introduced the density fields 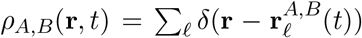. (The sum runs over the *N_A,B_* particles of species A or B whose positions are noted 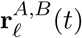.)

To account for the collective behaviors observed in experiments, we constructed coarse-grained dynamics describing the temporal evolution of the local cell densities of each species on time-scales much larger than the durations of the run and tumble phases, but shorter than the typical division time. These dynamical equations for the density fields *ρ_A,B_*(r,t) are given by (see SI):

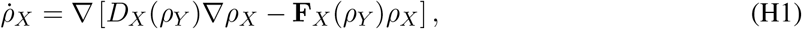

with (*X,Y*) ∈ {(*A,B*), (*B,A*)} and

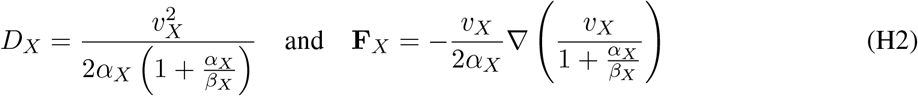

## 2. Linear stability analysis

We first performed a linear stability analysis of Eq. (H1) around homogeneous profiles 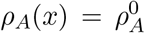 and 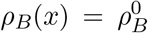. The Fourier modes of a fluctuation then evolve as 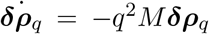 where 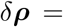 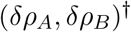 and *M* is a 2 × 2 matrix whose entries read

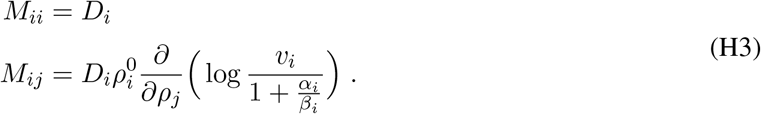

where *D*_*i*_ is defined in Eq. (H2). Note that the density-dependent parameters *v*_*i*_, *α*_*i*_ and *β*_*i*_ are evaluated at 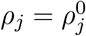 The condition for *M* to have an eigenvalue with negative real part, hence leading to an instability, is given by:

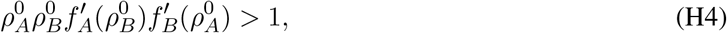

where we have defined the functions *f*_*A*_ and *f*_*B*_ as:

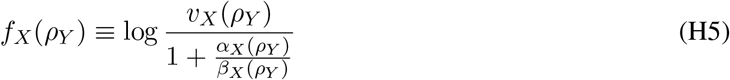

Note that in the presence of such an instability, one needs to consider higher order gradients, neglected in Eq. (H1), to stabilize smooth interfaces (see SI). These terms take the form −*κ*Δ^2^*ρ*_*X*_ and are phenomenologically included in Eq. (1) in the main text.

We note that *f*_*X*_ increases when *v*_*X*_ or *β*_*X*_ (resp. *α*_*X*_) are increasing (resp. decreasing) functions of *ρ*_*Y*_, corresponding to an enhancement of motility, whereas it decreases in the opposite case corresponding to an inhibition of motility. The sign of its derivative therefore determines whether the motility of species *X* is activated 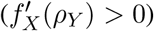 or inhibited 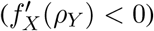 by species *Y*. The instability criterion (H4) requires the *product* 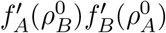 to be *positive*; it can be satisfied in two ways:

i. 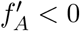 and 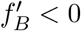: this corresponds to mutual inhibition;
ii. 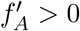 and 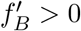: this corresponds to mutual activation.

Note that condition (H4) predicts no instability when 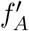 and 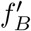 have opposite signs which we confirmed by simulations: systems of two species with mixed inhibition and activation of motilities do not undergo pattern formation.

Let us now explain why mutual inhibition of motility leads to aggregation of the two species whereas mutual activation leads to demixing. We consider the eigenvector 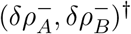 corresponding to the unstable eigenvalue *σ*^−^ < 0, whose properties are informative about how a small perturbation evolves in time. The unstable eigenvalue is given by

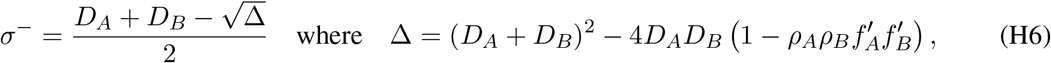

from which a corresponding eigenvector can be easily derived:

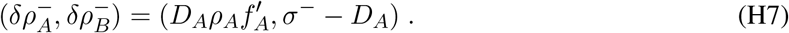

We observe that the sign of 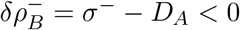 is fixed, whereas the sign of 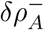 given by the sign of 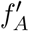 We detail the two possible cases, illustrated in Supplementary Fig. 10:

i. Mutual inhibition of motility corresponds to 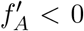 and 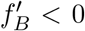 and leads to 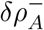 and 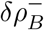 of the same sign. The densities of the two species thus grow (or decrease) in the same regions of space, leading to colocalization;
ii. Mutual activation of motility corresponds to 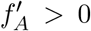 and 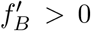 and leads to 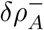 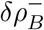 and of opposite signs. The density of one species thus grows (resp. decreases) in regions of space where the density of the conjugate species decreases (resp. grows), leading to the demixing of the two species.

So far we have considered the most general case, in which all motility parameters (the swimming speed *v*, the tumbling rate *α*, and the swimming rate *β*) of one species can depend on the density of the conjugate species. In the experiments, the parameter controlled by the QS interactions is the expression of *cheZ*, enhanced in the case of mutual activation and repressed in the case of mutual inhibition. Since CheZ controls the rate at which bacteria resumes swimming [17–20], we used density-dependent *β*_*A,B*_(*ρ*_*B,A*_) in our simulations, but kept *α*_*A,B*_ and *v*_*A,B*_ constants. Our model thus show these interactions to be sufficient to account for the types of patterns seen in experiments: the two species are colocalized in mutually inhibiting systems whereas they segregate in mutually activating ones.

## 3. Population dynamics

In our experiments, the full development of the patterns takes almost a day of incubation. On such time scales, two competing processes take place. First, the mechanism described above promote an unlimited coarsening of the patterns emerging from the linear instability. Then, the division of bacteria promote a local cell density equal to the uniform carrying capacity set by the medium. To determine the outcome of this competition, we added a logistic term to the dynamics (H1) to model bacterial growth and starvation, as frequently done in the literature [7]. Starting from a central inoculum, our simulations show a radial spreading of the colony followed by the formation of patterns whose growth saturates, leading to concentric rings of finite widths. The latter respects the local organization imposed by the instability mechanism: mutual activation of motility leads to rings of alternate compositions whereas mutual inhibition leads to colocalization of the two species within each high density ring.

## Appendix I: Simulations

Simulations of the dynamics (1) are performed with pseudo-spectral methods for the calculation of the spatial derivatives and semi-implicit Euler method for the time integration.

